# Individual variation in breeding phenology and postnatal development in northern bats (*Eptesicus nilssonii*)

**DOI:** 10.1101/2024.05.22.595341

**Authors:** Mari Aas Fjelldal, Jeroen van der Kooij

**Affiliations:** Finnish Museum of Natural History, University of Helsinki, Helsinki, Finland; Nature Education, Research and Consultancy van der Kooij, Rudsteinveien 67, Slattum, NO-1480, Norway

**Keywords:** Reproduction, gestation, parturition, neonatal, Chiroptera

## Abstract

Bats inhabiting northern latitudes are faced with short reproductive seasons during which they must produce and rear pups before fattening up in time to survive the winter hibernation. Therefore, the timing of parturition has considerable impacts on future fitness prospects for mother and pup; however, little is known about individual variation in breeding phenology and its consequences for postnatal development within bat populations. Here, we studied the phenology of breeding in *Eptesicus nilssonii* across seven years using data collected by day-to-day monitoring of a breeding colony in Norway (60.1°N) for which the identity and age of each mother (N = 8) and pup (N = 28) were known. By applying mixed-effect models, we found that arrival at the colony was largely dependent on late spring temperatures for all females, but that there were strong and consistent individual differences in arrival time across years. Females generally arrived ∼31.6 days (± 0.8 SE) before giving birth but could delay the timing of parturition by leaving the colony during early gestation if faced with poor weather conditions. However, females arriving late expressed shorter gestations, and pups born later in the season were born smaller but had higher growth rates during the most rapid growth period (<10 days old). The within-individual effects suggest that the higher growth rates could be due to mothers compensating for late parturition rather than by improved food availability. Date of parturition did not influence adult body size in pups. Pups became volant at the earliest only 13.1 days after birth (mean: 15.3 ± 1.6 SD) and approached adult flight patterns during their first flight week. Our unique results suggest that *E. nilssonii* is highly adapted to a short breeding season and is able to buffer unfavourable weather conditions to avoid slowing pup development, although the mechanistic drivers remain unknown.

## 1 Introduction

For many species in seasonal environments, spring and summer are a critical time of year when resource abundance and warmer thermal environments allow individuals to invest in reproduction (Fretwell 1972). However, at northern latitudes the spring season can be inhospitable while the summer season is short. The timing and success of reproduction in high-latitude environments therefore tend to fluctuate annually and largely depend on spring and summer conditions (e.g. Linton and Macdonald 2018, Fjelldal, et al. 2020).

Female bats at northern latitudes follow a reproductive cycle where they mate in autumn, store sperm in the uterus throughout the winter hibernation, initiate fertilization and gestation in spring, before reaching parturition in late spring or early summer (Racey 1982). Because shorter summer seasons limit the time available for fostering pups and rebuilding fat reserves for the upcoming winter hibernation, reproductive females and their offspring generally benefit from earlier parturition dates in seasonal environments (Ransome and McOwat 1994, Frick, et al. 2010, Linton and Macdonald 2018). However, females that initiate gestation early may obtain earlier parturition dates, but risk giving birth and entering the energetically costly lactation period before food availability has stabilised (Rydell 1989). Pregnant bats therefore need to balance the risk of facing an energetic mismatch by giving birth early against the long-term fitness costs associated with giving birth late.

The trade-offs associated with the timing of parturition is naturally strongly dependent on interannual spring conditions, where cold, wet and windy weather conditions delay the breeding phenology in bats (e.g. Linton and Macdonald 2018, Sunga, et al. 2023). During poor environmental conditions in spring, pregnant females can reduce energetic costs by employing torpor, although this pauses the foetal development and prolongs gestation (Racey and Swift 1981, Willis, et al. 2006). The influence of spring conditions on the timing of parturition in bats is therefore likely caused by a combination of delayed emergence from hibernation sites and increased use of daily torpor during the gestation period. However, after accounting for environmental effects we still lack knowledge on differences in individual reproductive strategies, or ‘personalities’ (Dingemanse, et al. 2010), in insectivorous bats, which potentially exist along a behavioural continuum with two extremes: initiate gestation early to prolong the breeding season and cope with a more stochastic environment, or initiate gestation late during more stable conditions and cope with a limited breeding season and reduced potential to build up fat reserves for the winter.

For bats belonging to the family of Vespertilionidae, which is the largest family within the order of Chiroptera, the pups are born large (neonatal forearm length ranging from 25% to 43% of maternal forearm-length; Kurta and Kunz (1987)) and express rapid early development rates compared to what would be expected from the low annual recruitment and high longevity (Barclay, et al. 2003). In northern latitude environments with short breeding seasons, mammals are expected to adopt fast postnatal growth rates to enhance survival chances across the approaching winter season (Boyce 1979). Observations of postnatal development in bats across latitudes align with these predictions, where a strong selection pressure for faster postnatal growth rates was found in pups born in temperate areas compared to tropical climates (Kunz and Stern 1995). Breeding colonies of northern bats (*Eptesicus nilssonii*) are found as far north as 69ºN (Rydell, et al. 1994, Frafjord 2013, Suominen, et al. 2022) which expose them to prolonged periods of continuous daylight under the midnight sun. Being the northernmost breeding bat species in the world, *E. nilssonii* can be expected to possess behavioural, physiological and/or life-history adaptations that allow them to survive and reproduce at latitudes that are too challenging for most other bat species. However, the exact mechanisms allowing this species to overcome the energetic bottlenecks of costly breeding in such areas are still unknown.

Here, we explore unique individual-based data across seven consecutive years (2017 to 2023) of a small maternity colony of *E. nilssonii* in Norway. Such long-term individual-based data are scarce within the order of Chiroptera (but see Linton and Macdonald 2018), and detailed day-to-day monitoring of mothers and pups have not previously been recorded in free-ranging bats. In this study we investigate individual level breeding phenology and postnatal development in a colony faced with the challenges of living in a high-latitude (60ºN) environment. We hypothesize that breeding phenology is negatively impacted by poorer spring conditions, and that unfavourable weather conditions (i.e., low environmental temperatures, heavy rain, strong winds, etc.) during the early growth period have negative impacts on pup development, in accordance with observations from other studies (e.g. Hoying and Kunz 1998, Linton and Macdonald 2018). Additionally, we explore maternal differences across years and potential adaptations to the challenges of breeding in a high-latitude environment.

## 2 Materials and Methods

### 2.1 Data collection

Data was collected during seven consecutive breeding seasons between 2017-2023 in Nittedal, Norway (60.11°N, 10.85°E). A small maternity colony consisting of a total of eight reproductive female *E. nilssonii* with known identities and ages (see section 2.1.1) roosted in a SW orientated/exposed wooden bat box on a garage wall 1.5 m above the ground. The box was equipped with a heating device (135-Watt heater with thermostat) at the top; however, this was turned off on very warm days (air temperatures > ∼25ºC) to avoid overheating. A camera (D-link DSC-2670L) was installed in the box before the breeding season of 2019, allowing real-time observations through the software mydlink (version 2.11.0, D-Link Corporation) in addition to motion-triggered video-recordings. Infrared sensors (ChiroTec Tricorder 9600) were installed at the entrance of the box before the breeding season of 2020 and recorded the timing of each emergence and return at night.

The bat box was manually checked twice per day throughout the breeding seasons; once in the morning to check for the presence of each individual and to (occasionally; five to 15 times per mother per season) take biometric measurements of each adult (weights to nearest 0.1 g using a digital mini-weight and forearm-length to nearest 0.1 mm using dial callipers); and once in the evening after pups were born in order to take daily biometric measures and photographs of their development. When approaching the time of parturition, the box was occasionally checked also during mid-day to get accurate timing of each birth and to get forearm measurements of newborn pups, although suckling pups never were separated from their mothers. The evening measurements of pups were conducted after the females had left the box to forage at night and were always accomplished as a fast as possible (finished within five to ten minutes per night). Because most of the females in the colony had been hand-raised as pups (see section 2.1.1) and therefore were used to eating mealworms, a few mealworms (∼1g per bat) and fresh water were supplied after every morning count as compensation for the disturbance; however, the females that had been born in the box to later return as part of the colony did not eat mealworms (verified through camera-observations).

Across the study period the eight individual females gave birth to a total of 28 viable pups and one stillborn Siamese twin. We recorded 1-7 breeding seasons per female (average 3.3 ± 2.5 *SD*). Four females gave birth for the first time as one-year olds, while four females gave birth for the first time as two-year olds.

#### 2.1.1 Individual identification

Of the eight females, six had been brought in as abandoned pups and were hand-reared until they reached adult size, upon which they were released in the bat box, but were still supplied with additional mealworms until they left the box in autumn for the upcoming winter hibernation. In consecutive years, these six hand-reared females returned to the bat box in spring to give birth. The remaining two females in the colony were daughters born in the box that returned the following summers as part of the maternity colony. The identity of each bat was confirmed at the beginning of each breeding season by photographing their wings, using an external flash as a backlight, as patterns of collagen–elastin bundles can be used as an individual identifier (Amelon, et al. 2017). We confirmed that this was a successful method for identification in our small maternity colony by investigating cross-year wing-photos taken of the two oldest females, which had been ringed as juveniles. Given the small number of bats present each year, the identification throughout the summer was afterwards made through measurements of forearm-length and observations of individual morphological characteristics and was re-confirmed by wing-photos taken during the season. Except for on one occasion, pups were always born with a few days intervals and were therefore easily identified by their postnatal development stage until they reached adult size, upon which forearm-measurements, visual morphological characteristics and wing-photos were used to confirm their identities.

#### 2.1.2 Phenology

We recorded arrival day and daily presence of each female in the colony, timing of parturition, the timing of the pups being volant (observed either on camera or by registering absence during the evening box check), and the timing when adults and juveniles left the colony in late summer.

#### 2.1.2 Meteorological data

We downloaded weather data with 10-minute intervals from the meteorological stations SN4460 (temperature, wind and rain) and SN18700 (barometric pressure) through the Norwegian Centre for Climate Services (NCCS). Sunset and sunrise data were obtained through the Time and Date webpage (Timeanddate.com).

### 2.2 Data analyses

We performed all analyses in R (version 4.3.1). Mixed models (LMMs or GLMMs) were fitted using the *lmer* or *glmer* functions from the lme4 package (Bates, et al. 2015). We based our model selection on Akaike information criterion corrected for small sample sizes (AICc) and model weights using the *dredge* function from the MuMIn package (Barton and Barton 2015). We performed initial smaller models in cases where several versions of the same variable were of interest (i.e. mean, maximum and minimum temperature values for day-time or night-time) to decide which variable to include in the global models, based on AICc values. To disentangle within-versus among-individual effects of explanatory variables in the final models (i.e. after model selection), we applied the mean-centering methods described in van de Pol and Wright (2009) for distinguishing the effects of within-individual plasticity versus among-individual differences.

#### 2.1.2 Arrival phenology and gestation period

We fitted LMMs to test which spring weather conditions that best explained the variation in the arrival date (day in relation to 1^st^ May) to the colony, with year and female ID included as random effects. The fixed effects tested were the onset of spring (the ordinal day when the 10 day smoothed daily temperature is above 0°C for at least ten days (Fjelldal, et al. 2020)), or either mean, min or max temperature conditions for three selected time periods: from 1^st^ April to 30^th^ April, from 15^th^ April to 15^th^ May, and 1^st^ May until 31^th^ of May. We chose to limit the number of time periods tested due to our small sample size (N_obs_ = 28). Total rainfall for each period was also included as a fixed effect in each model. We then performed model selections on each global model, and the AICc of each of the highest ranked models were compared to see which spring weather condition that best explained arrival time.

During the gestation period pregnant females occasionally left the bat box to return a few days later. To test which weather conditions affected these periods of absence, we treated individual presence in the colony during the gestation period as a binary response (present = 1, absent = 0) in GLMMs (family = “binomial”, link = “logit”). Maximum air temperature and total precipitation on the day before the observation were included as fixed effects. We also categorized each presence observation into ‘early gestation’ (≥ 15 days left until parturition) or ‘late gestation’ (< 15 days left until parturition) and tested temperature and rainfall in interaction with gestation stage. We chose to categorize the time left until parturition due to high correlations with the temperature variables, and due to the observations of females being absent were made either very early in the gestation period (from 24.5 to 36.7 days left until parturition, mean = 30.9 days ± 3.2 *SD*, N_obs_ = 13) or very late (from 0 to 10.7 days left until parturition, mean = 6.7 days ± 0.9 *SD*, N_obs_ = 38). Year and female ID were included as random effects.

To assess the effect of the missing periods on the gestation duration we fitted a LMM that included the number of days of absence from the colony in the early and in the late gestation period as fixed effects, and year and female ID as random effects.

#### 2.2.2 Postnatal growth

Variation in size at birth (body weight and forearm-length) was tested with global LMMs that included age at measurement, sex, age of the mother, whether the mother ate offered mealworms or not, gestation time, birth date in relation to 1^st^ of June, total rainfall and mean temperature the last three weeks before parturition as fixed effects, while year and female ID were included as random effects.

We constructed logistic, Gompertz and von Bertalanffy growth models on forearm length and body mass and compared AICc for the best fit. However, each growth model consists of a rapid, approximately linear growth period before the growth declines. To determine the duration of the most rapid growth period we performed breakpoint analyses from the segmented package (Muggeo 2008). Finally, we tested the effects on the most rapid growth period by constructing global LMMs with individual growth rates during the most rapid growth period as response variables, and sex, age of the mother, whether the mother was eating offered mealworms or not, birth date in relation to 1^st^ of June, total rainfall and minimum air temperatures at night during the most rapid growth period as fixed effects, with year and female ID included as random effects.

Forearm-length and body mass variation in pups approaching adult size were explored by fitting LMMs with the individual largest recorded forearm-length or heaviest body weight (before 3 weeks of age, as several pups were not measured after this point in time) as response variables. Sex, age of the mother, whether the mother ate offered mealworms or not, birth date in relation to 1^st^ of June, total rainfall and mean temperature the three first weeks after birth were included as fixed effects, and year and female ID were included as random effects.

## 3. Results

### 3.1 Breeding phenology and gestation period

The females generally arrived at the colony in May and gave birth to their pups in June (Table 1). The model that best explained the variation in arrival date in spring included a negative effect of mean temperature between 15^th^ April to 15^th^ of May, which we confirmed was driven by within-subject effects (Table 2a; N_obs_ = 28). However, although each female showed similar responses of arriving ∼4 days earlier per one degree increase in mean temperature conditions (Table 2a; Fig. 1a), there were strong individual differences in arrival time. Figure 1b illustrates our observations of how each female arrived at certain time intervals when compared to a reference female that was present across all seven breeding seasons, showing that some females arrived generally early while others arrived late compared to each other.

**Table 1:** Phenological and developmental observations made across the study period.

| Observations | Dates | Time since birth (mean ± sd) | N<sub>obs</sub> |
| --- | --- | --- | --- |
| Females arrive in the colony | 30/04 to 04/06 | -23.6 to -40.4 days (-33.5 ± 5.4) | 28 |
| Parturition | 07/06 to 27/06 |  | 28 |
| Last obs. of umb. cord* | 07/06 to 28/06 | 1.7 to 26.3 hours (median = 4.6) | 24 |
| Pups open eyes | 09/06 to 01/07 | 1.6 to 5.4 days (4.1 ± 0.9) | 26 |
| Pups' first flight | 24/06 to 13/07 | 13.1 to 19.4 days (15.3 ± 1.6) | 25 |
| Mothers leave the colony | 05/07 to 03/08 | 14.1 to 46.2 days (32.7 ± 8.1) | 21 |
| Pups leave the colony | 05/07 to 31/07 | 14.1 to 40.4 days (30.6 ± 7.0) | 18 |

\* Last observation where the umbilical cord was still attached

**Table 2:**
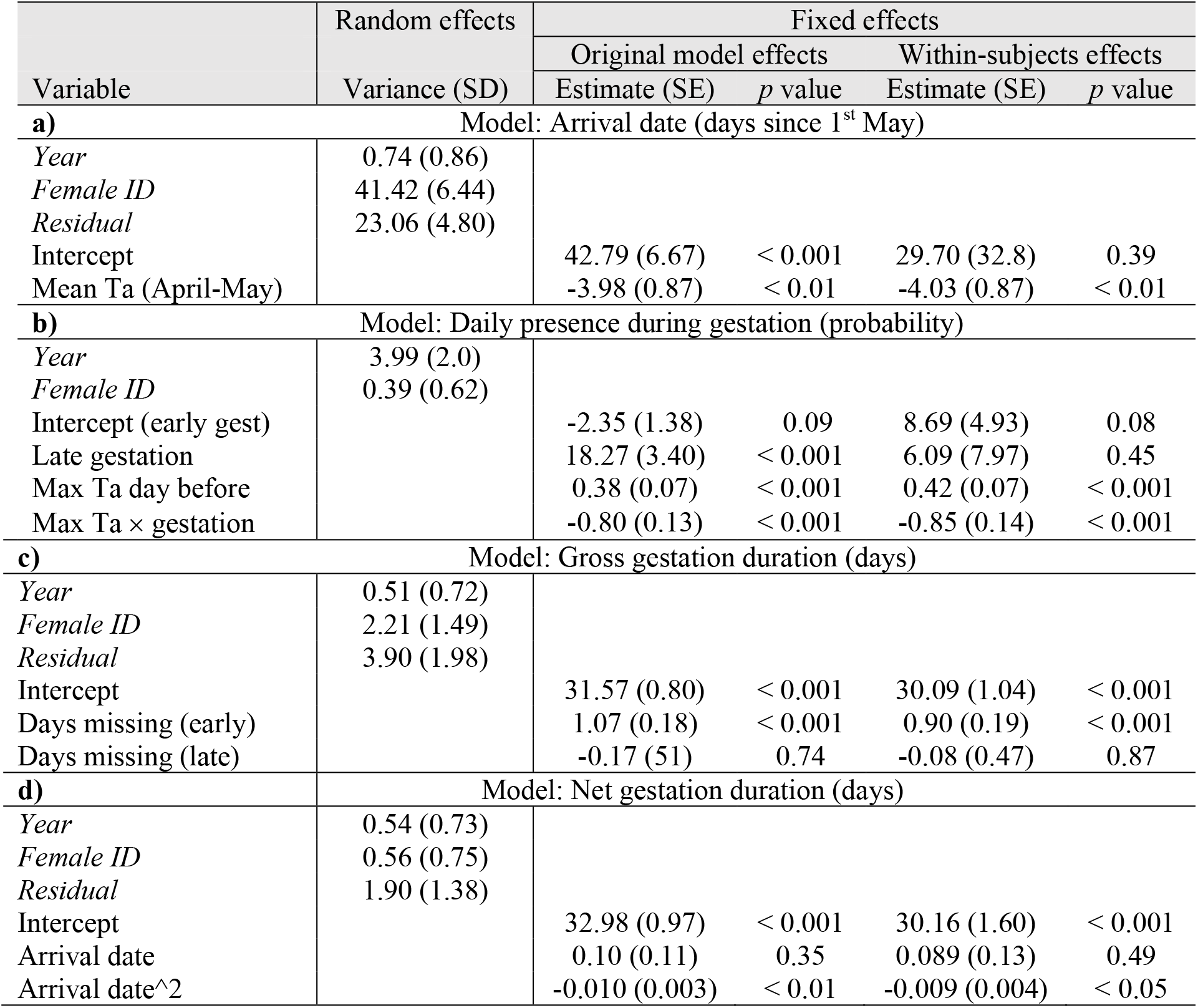
Model results presenting original (undecomposed) and decomposed within-subject effects from the final best-fit models (after model selection) explaining: **a)** arrival date; **b)** presence in colony during gestation period; **c**) gross gestation period duration; and **d)** net gestation period duration.

**Figure 1:**
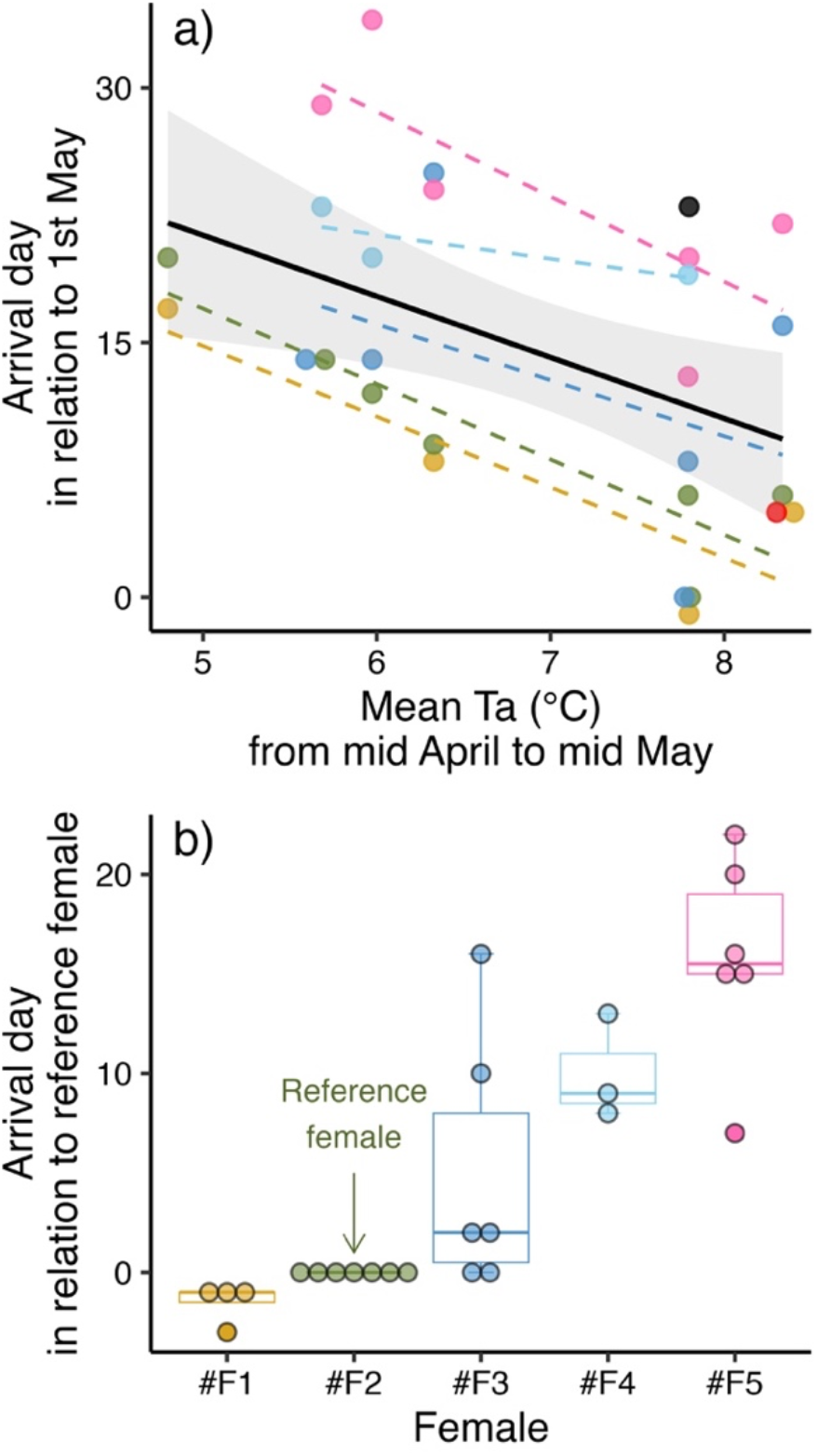
Variation in arrival day at the breeding colony. **a)** The effect of increasing mean temperature conditions between 15^th^ April to 15^th^ of May leading to earlier arrival dates in spring, with the black regression line showing the overall effect while different colours indicate different females. **b)** Timing of arrival for each female when compared to a reference female (*#F2*). *Note*: Three females were only present for one breeding season and are therefore not shown with individual regression lines or boxplots.

Daily presence probability in the colony during the gestation period was best explained by an interaction effect between maximum temperature the day before and the gestation stage (‘early’ or ‘late’) (Table 2b; N_obs_ = 874). Colder maximum temperatures led to an increasing risk of females leaving the colony the following night in the early gestation period, while warmer maximum temperatures increased the risk of leaving the colony in the late gestation period (Fig. 2a). The effects were confirmed to be driven by within-subject effects (Table 2b).

**Figure 2:**
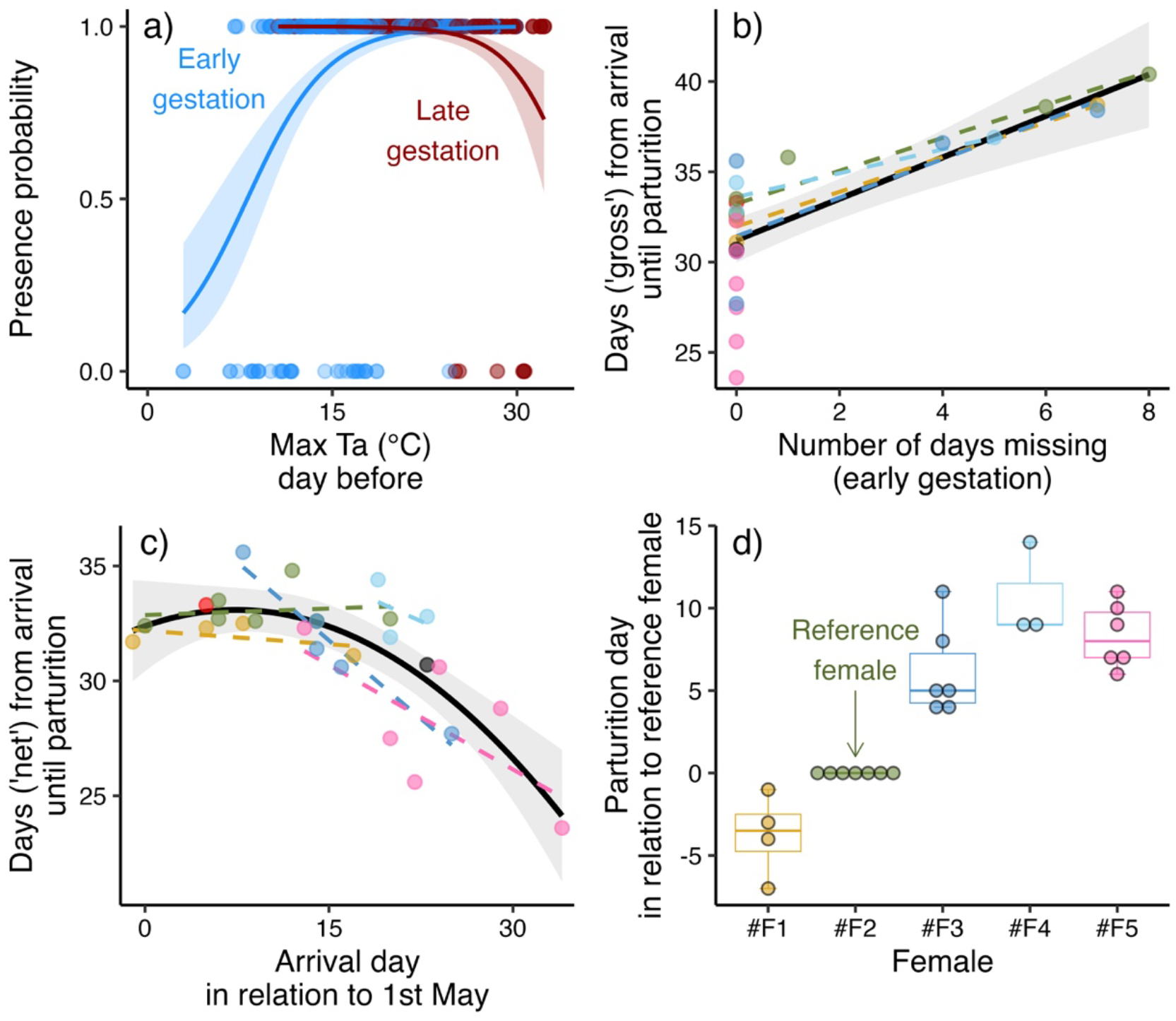
Gestation period dynamics. **a)** The probability of a female being present in the colony in relation to maximum temperature the day before, where colder temperatures decreased the presence probability during the early gestation period (blue), while warmer temperatures decreased the probability during the late gestation period (red). **b)** The positive effect of absent periods during the early gestation period, with the black regression line showing the overall effect while different colours indicate different females. **c)** Quadratic effect of arrival date on the net gestation time (subtracting time absent during early gestation), showing that later arrival dates were associated with shorter gestation time. The regression lines per female are shown as linear effects due to few datapoints. **d)** Timing of parturition for each female when compared to a reference female (*#F2*).

The periods of absence from the colony prolonged the estimated gestation period (31.6 days ± 0.8 *SE*; N_obs_ = 27) with the same number of days as the females were missing, but only during the early gestation period when the absence was caused by cold weather conditions (Fig. 2b). Females leaving the box late in the gestation period (due to overheating) did not prolong the gestation period (Table 2b; effects were confirmed to be within-subject effects). We therefore differentiate between ‘gross’ and ‘net’ gestation period: the gross gestation time refers to the time period from a female first arrived in the colony until giving birth, which lasted between 23.6 to 40.4 days. The net time subtracts the days females were absent during the early gestation period, lasting from 23.6 to 35.6 days. The net gestation time was shortened when females arrived later in spring (quadratic effect; Fig. 2c), which we confirmed was driven by within-subjects effects (Table 2c). We observed similar individual differences in the timing of parturition as we did for timing in arrival, although the female that usually arrived last (#F5) not always gave birth last (Fig. 2d) due to her overall shorter gestation periods (Fig. 2b and c).

### 3.2 Postnatal growth

Timing of parturition was recorded with high accuracy (from 0 to 6 hours uncertainty, mean: 2.0 hours ± 2.0 *SD*, N_ind_ = 27), except for on one occasion where a heavily pregnant female left the bat box due to high temperatures and returned to the colony three days later with her pup (estimated to have been between ∼1.5 to 2 days old based on forearm-length). A total of 7 female pups, 21 male pups and one stillborn Siamese twin was born during the study period (see Table S1 in Supplementary Materials). The last observations of umbilical cords being attached (sometimes with the placenta still connected, see Fig. S1 in the Supplementary Materials) was made from 1.7 to 26.3 hours after birth (median = 4.6 hours, N_ind_ = 24), although it was only once observed to be attached longer than 24 hours. Pups opened their eyes when they were between 1.6 and 5.4 days old (mean = 4.1 days ± 0.9 *SD*, N_ind_ = 26). The fur growth on abdomen and back during the most rapid growth period (see section 3.2.2) is illustrated in Figure 3.

**Figure 3:**
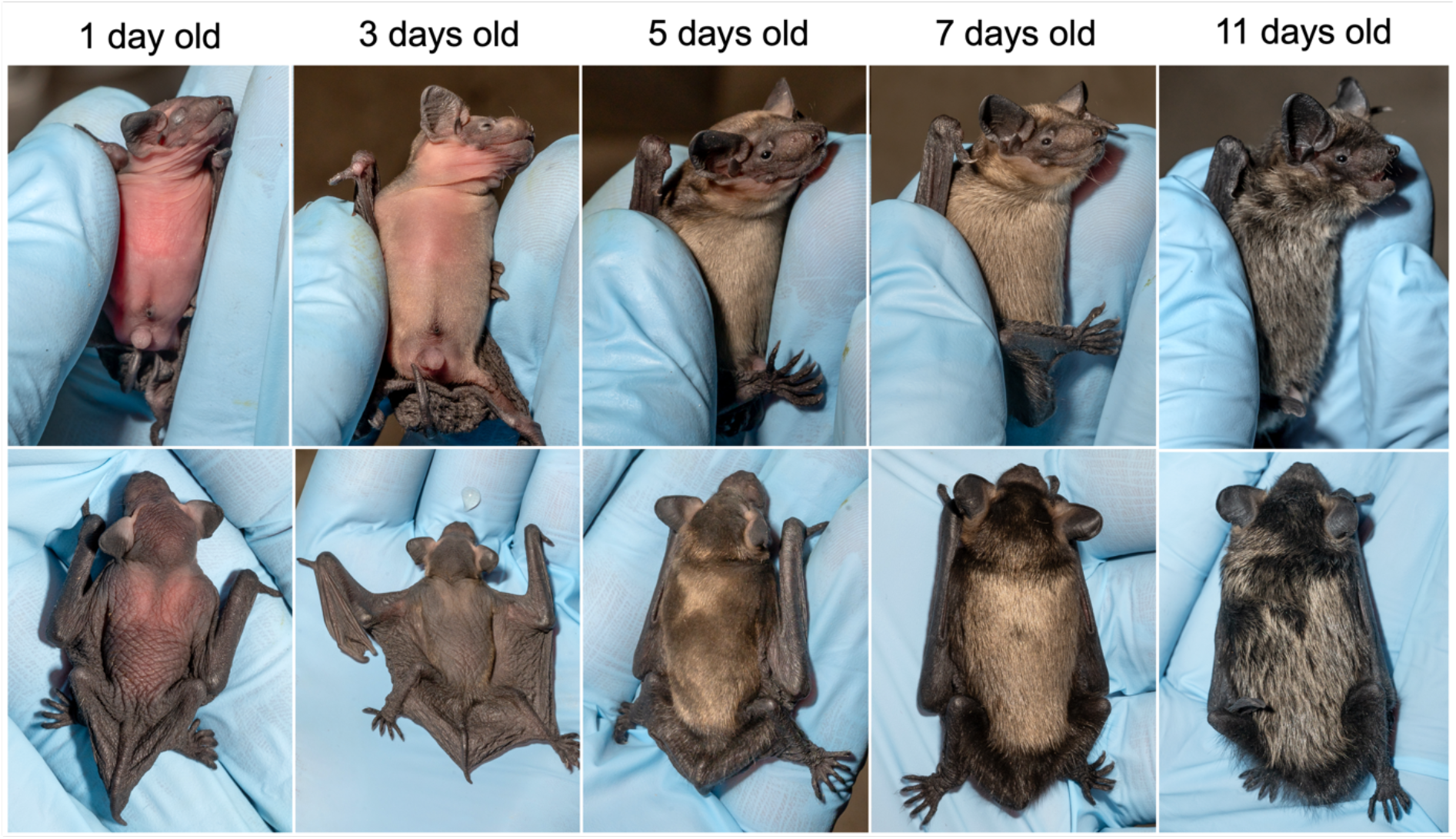
Morphologic development in a male northern bat pup during the first 11 days, demonstrating the gradual fur growth on the abdomen and on the back. The ages are exact (± 2 hours). Photos by Jeroen van der Kooij.

#### 3.2.1 Size at birth

The estimated body mass at birth when accounting for age at measurement was 2.2 g (± 0.17 *SE*, N_ind_ = 21, max age at measurement = 1.5 days), where newborn pups on average had body masses corresponding to 25.8% (± 0.06 *SD*, N_ind_ = 12) of their mothers’ mean postpartum body masses (each female was measured multiple times throughout the lactation period). Age at the time of measurements was the only variable explaining the variation observed in body mass (estimate: 0.047 g/hour ± 0.007 *SE*, p < 0.001). The estimated forearm-length at birth in northern bat pups when accounting for age at measurement was 15.7 mm (± 0.39 *SE*, N_ind_ = 25, max age at measurement = 0.7 days), with newborn pups on average having forearm-lengths corresponding to 41% (± 0.03 *SD*, N_ind_ = 16) of their mothers forearms. The model that best explained variation in forearm length at birth included a negative effect of birth date and a negative effect of total rainfall during the three last weeks before parturition (Table 3a; N_ind_ = 25). However, rainfall was heavily negatively correlated with mean temperatures (cor: -0.82, p < 0.001), and we can therefore understand increasing rainfall values as colder and wetter weather conditions. Later parturition dates and wetter/colder weather conditions the weeks before parturition thus resulted in smaller sized pups at birth (Fig. S2 in the Supplementary Materials), which we confirmed were driven by within-subjects effects (Table 3a).

**Table 3:**
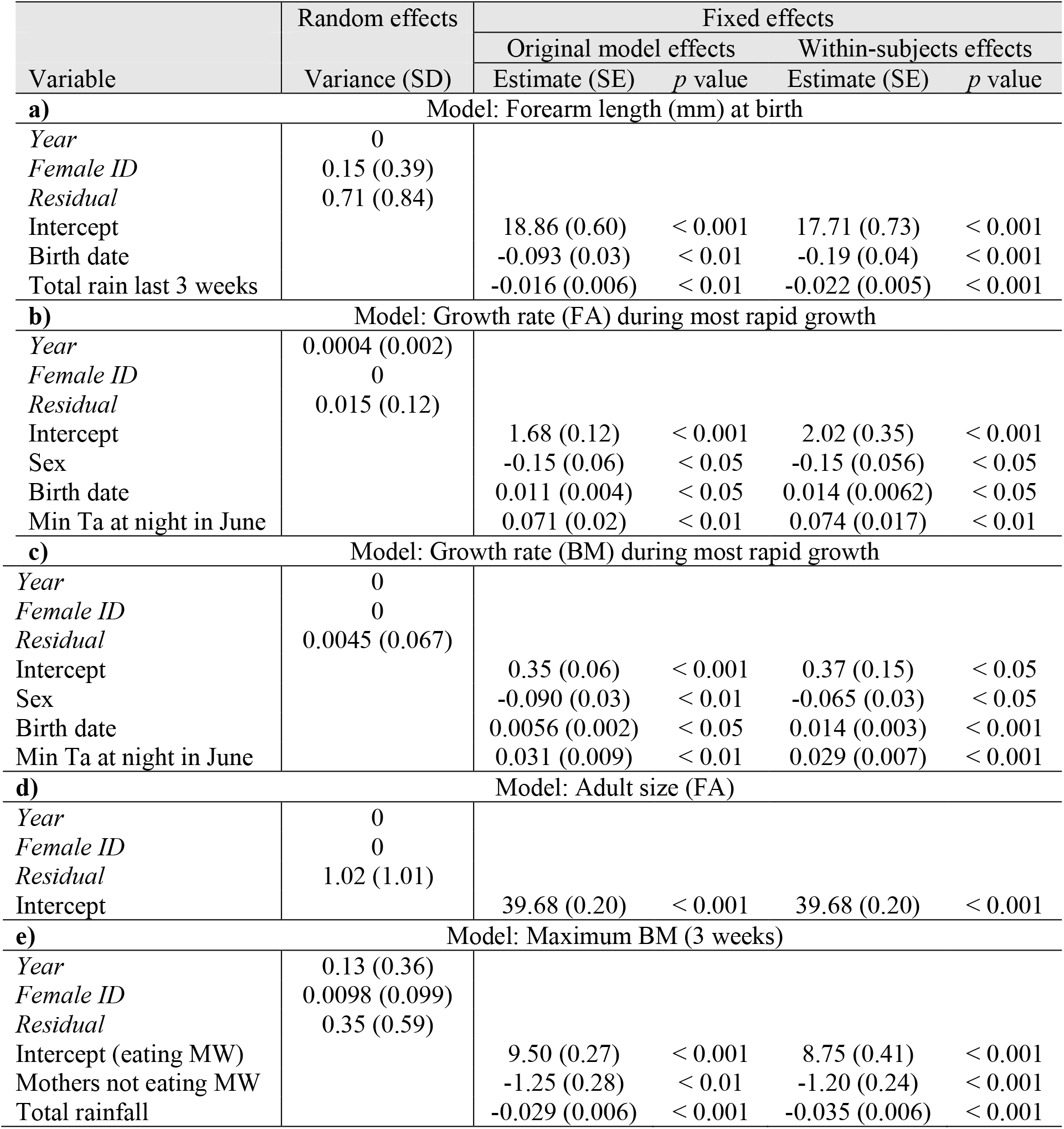
Model results presenting original (undecomposed) and decomposed within-subject effects from the final best-fit models (after model selection) explaining: **a)** forearm length at birth; **b)** forearm growth rate during the most rapid growth period; **c**) gestation period duration; **d)** forearm length at birth; **e)** adult forearm length; and **f)** maximum body mass during first three weeks (*note*: MW denotes ‘mealworms’).

| Variable | Random effects<br>Variance (SD) | Fixed effects |  |  |  |
| --- | --- | --- | --- | --- | --- |
|  |  | Original model effects |  | Within-subjects effects |  |
|  |  | Estimate (SE) | *p* value | Estimate (SE) | *p* value |
| **a)** Model: Forearm length (mm) at birth |  |  |  |  |  |
| *Year* | 0 |  |  |  |  |
| *Female ID* | 0.15 (0.39) |  |  |  |  |
| *Residual* | 0.71 (0.84) |  |  |  |  |
| Intercept |  | 18.86 (0.60) | < 0.001 | 17.71 (0.73) | < 0.001 |
| Birth date |  | -0.093 (0.03) | < 0.01 | -0.19 (0.04) | < 0.001 |
| Total rain last 3 weeks |  | -0.016 (0.006) | < 0.01 | -0.022 (0.005) | < 0.001 |
| **b)** Model: Growth rate (FA) during most rapid growth |  |  |  |  |  |
| *Year* | 0.0004 (0.002) |  |  |  |  |
| *Female ID* | 0 |  |  |  |  |
| *Residual* | 0.015 (0.12) |  |  |  |  |
| Intercept |  | 1.68 (0.12) | < 0.001 | 2.02 (0.35) | < 0.001 |
| Sex |  | -0.15 (0.06) | < 0.05 | -0.15 (0.056) | < 0.05 |
| Birth date |  | 0.011 (0.004) | < 0.05 | 0.014 (0.0062) | < 0.05 |
| Min Ta at night in June |  | 0.071 (0.02) | < 0.01 | 0.074 (0.017) | < 0.01 |
| **c)** Model: Growth rate (BM) during most rapid growth |  |  |  |  |  |
| *Year* | 0 |  |  |  |  |
| *Female ID* | 0 |  |  |  |  |
| *Residual* | 0.0045 (0.067) |  |  |  |  |
| Intercept |  | 0.35 (0.06) | < 0.001 | 0.37 (0.15) | < 0.05 |
| Sex |  | -0.090 (0.03) | < 0.01 | -0.065 (0.03) | < 0.05 |
| Birth date |  | 0.0056 (0.002) | < 0.05 | 0.014 (0.003) | < 0.001 |
| Min Ta at night in June |  | 0.031 (0.009) | < 0.01 | 0.029 (0.007) | < 0.001 |
| **d)** Model: Adult size (FA) |  |  |  |  |  |
| *Year* | 0 |  |  |  |  |
| *Female ID* | 0 |  |  |  |  |
| *Residual* | 1.02 (1.01) |  |  |  |  |
| Intercept |  | 39.68 (0.20) | < 0.001 | 39.68 (0.20) | < 0.001 |
| **e)** Model: Maximum BM (3 weeks) |  |  |  |  |  |
| *Year* | 0.13 (0.36) |  |  |  |  |
| *Female ID* | 0.0098 (0.099) |  |  |  |  |
| *Residual* | 0.35 (0.59) |  |  |  |  |
| Intercept (eating MW) |  | 9.50 (0.27) | < 0.001 | 8.75 (0.41) | < 0.001 |
| Mothers not eating MW |  | -1.25 (0.28) | < 0.01 | -1.20 (0.24) | < 0.001 |
| Total rainfall |  | -0.029 (0.006) | < 0.001 | -0.035 (0.006) | < 0.001 |

#### 3.2.2 Growth curves and most rapid growth period

The logistic model best explained the growth in forearm length (N_obs_ = 450, N_ind_ = 26) with equation: FA (mm) = 39.945 / 1 + e^-0.253 * (days old – 1.925)^ (Fig. 4a), while the von Bertalanffy model best explained body mass increase (N_obs_ = 424, N_ind_ = 26) with equation: BM (g) = 8.740 * (1 - e^-0.144 * (days old + 1.862)^) (Fig. 4b). Model results can be found in Table S2 in the Supplementary Materials, along with figures of individual growth for forearm length (Fig. S3) and body mass (Fig. S4) in each pup.

**Figure 4:**
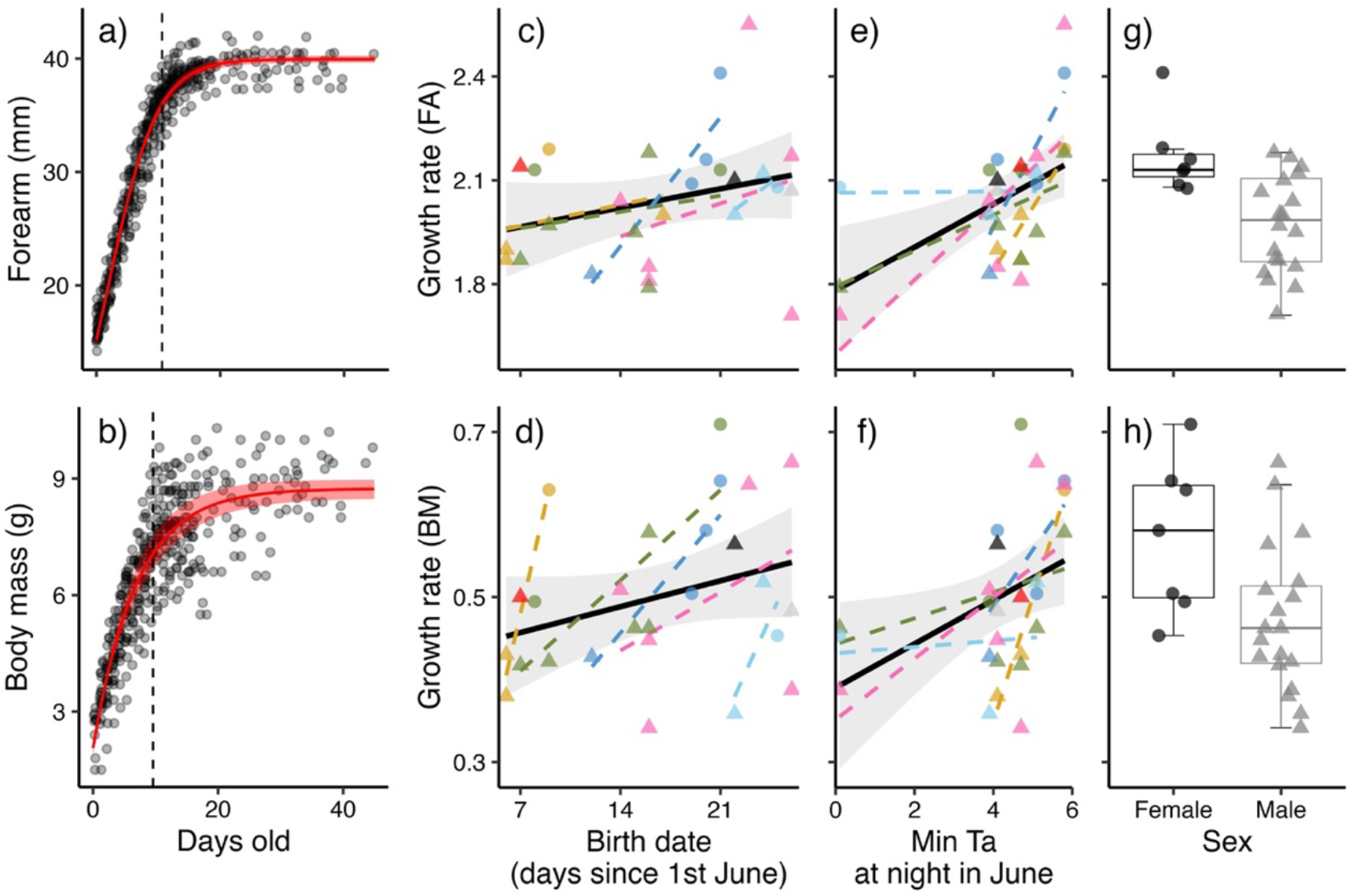
Postnatal growth curves and effects on individual growth rates during the most intense growth period in northern bat pups. **a)** Forearm length in relation to age with estimates from the logistic growth model shown as a red line (confidence intervals included as shaded light red). **b)** Body mass in relation to age with estimates from the von Bertalanffy growth model shown as a red line. Dashed lines indicate the breakpoint estimates, marking the end of the most rapid growth period. The variation observed in individual growth rates during this period was explained by **c-d)** a positive effect of birth date, **e-f)** a positive effect of minimum nightly temperature in June, and **g-h)** a sex effect, with female pups expressing higher growth rates than male pups. Different colours indicate different mothers, circles mark female pups and triangles mark male pups.

The breakpoint estimate indicating the end of the fastest growth period for growth in forearm length was at 10.6 days (± 0.13 *SE*) and for growth in body mass at 9.6 days (± 0.3 *SE*). During this period, starting at birth, the growth is approximately linear, and the model that best explained the variation in individual growth rates during the most rapid growth period (for both forearm and body mass) included sex, date of birth and minimum nightly temperature conditions in June (N_obs_ = 26). For both forearm length and body mass, female pups had higher growth rates than males, and later birth dates and warmer nights in June led to higher growth rates, with all effects being driven mainly by within-subjects effects (Fig. 4c-h; Table 3b & c). The within-subject effect was considerably stronger than the original non-decomposed effect for the effect of birth date on body mass growth rate (Table 3c), as seen in the steeper individual regression lines per mother in Fig. 4d.

#### 3.2.3 Adult body size

Forearm length for pups approaching adult size varied from 37.4 mm to 42.0 mm (mean: 39.7 mm ± 1.03 *SD*, N_ind_ = 26). None of the tested fixed effects explained the observed variation in adult forearm size, and the highest ranked model therefore only included intercept (Table 3d). Maximum individual body mass for pups being less than three weeks old ranged from 6.2 g to 10.3 g (mean: 8.3 g ± 1.1 *SD*, N_ind_ = 27). The variation in maximum body mass was best explained by a negative effect of total rainfall during the first three weeks after birth, and an effect of whether the mothers ate offered mealworms or not (see Methods section 2.1), with mealworm-eating mothers having heavier pups (Table 3e).

### 3.3 First flight week

The age at first flight ranged from pups being 13.1 days to 19.4 days old (mean age: 15.3 days ± 1.6 *SD*, N_ind_ = 25). On their first flight night, juveniles emerged between 48 minutes to 4 hours and 35 minutes after sunset (mean: 2.7 hours ± 1.2 SD, N_ind_ = 10), while adult individuals, for comparison, emerged between 0 minutes to 1 hour and 40 minutes after sunset (mean: 28.5 minutes ± 18.6 SD, N_obs_ = 38) on the same nights (Fig. 5a). Juveniles delayed their emergence time compared to adults also on their second flight night but gradually approached the emergence time observed in adults across the week after their first flight (Fig. 5a & Table S3 in the Supplementary Materials). Furthermore, the duration of the juvenile flights on the first flight night was considerably shorter compared to adults on the same nights, ranging from 1.3 minutes to 40.6 minutes in duration (mean: 15.3 minutes ± 15.8 *SD*, N_obs_ = 7) for juveniles and from 5 minutes to 252 minutes in duration (mean: 62.7 minutes ± 45.2 *SD*, N_obs_ = 46) for adults. On nights thereafter, the juvenile flights increased gradually in duration until they approached adult flight durations after about a week after their first flights (Fig. 5b & Table S4 in the Supplementary Materials).

**Figure 5:**
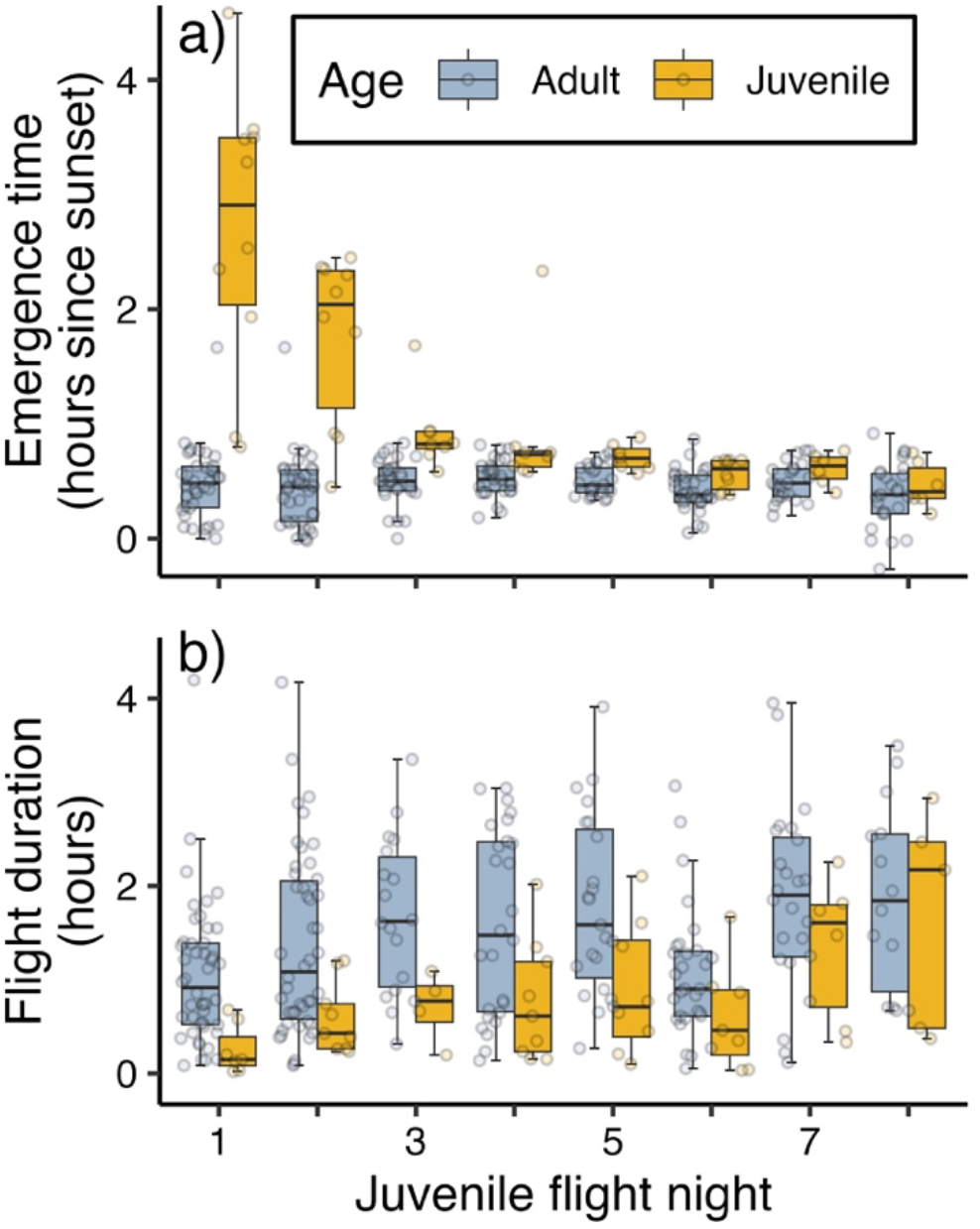
Juvenile flight patterns (yellow) compared to adults (blue) across the juveniles’ first flight nights in regards to **a)** emergence time (number of hours since sunset), and **b)** flight duration (hours).

### 3.4 Notes on mortality

One male pup died of unknown causes when he was eight days old, while one female pup never returned from her first flight at the age of 14 days. Of the six remaining female pups that left the colony after the breeding season, two never returned to the colony while three returned to the box after surviving their first winter. Two of these returning female pups produced pups of their own in the following years, while the third pup returned as an unreproductive one-year old in the final year of the study (Table S1). The last female pup was born during the final year of the study and can therefore not be evaluated.

## Discussion

Here, we present evidence of individual variation in breeding phenology in a high-latitude study population of *E. nilssonii*, with strong and consequent differences observed between females across years. Although late spring temperatures affected the arrival time of all females equally (i.e. similar individual plasticity), some females were consistently early while others were consistently late when compared to each other (i.e. variation in personalities (Dingemanse, et al. 2010)). These observations support the indication of among-individual differences observed in parturition dates for *Myotis lucifugus* in the study by Sunga, et al. (2023).

In temperate environments, poor spring conditions can drastically skew the breeding season and impose long-term detrimental fitness-consequences at the population level in bats (e.g. Frick, et al. 2010, Lučan, et al. 2013, Linton and Macdonald 2018). We found that at the individual level, reproductive *E. nilssonii* females attempted to compensate for such effects; bats arriving later at the colony reduced the duration of their gestation period, which aligns with the observations made in a captive population of *Nyctalus noctula* (Zukalova, et al. 2022). Also, while pups born later in the season were born smaller, they expressed higher growth rates than pups born earlier. This effect was substantially stronger as a ‘within-mother’ effect for the growth rate of body mass in pups (Fig. 4d, Table 3c), which suggests that the effect was caused mainly by females compensating for late parturition rather than by an improvement in food availability. Although higher growth rates in bat pups have been reported as a response to shorter breeding seasons across populations (Kunz and Stern 1995), this has never before been shown to apply within bat populations.

Weather conditions and food availability during spring and summer are already frequently reported as main drivers of intraspecific variation in breeding phenology (Rydell 1989, Ransome and McOwat 1994, Arlettaz, et al. 2001, Lučan, et al. 2013, Linton and Macdonald 2018, Matthäus, et al. 2023), reproductive success (Burles, et al. 2009, Lučan, et al. 2013, Linton and Macdonald 2018) and postnatal development in bats (Hoying and Kunz 1998, Koehler and Barclay 2000, Hood, et al. 2002, Mundinger, et al. 2021). In our study population, environmental conditions during spring and summer influenced both breeding phenology and postnatal development, despite the box being heated and a few mealworms being supplied daily. Lower temperatures and rainier conditions delayed arrival time to the colony, reduced daily presence probabilities in the bat box during early gestation, decreased forearm length in pups at birth, decreased daily postnatal growth rates, and resulted in lighter juvenile body mass in three-week-old pups. However, we did not detect any environmental, phenological, or maternal effects on the variation in final juvenile forearm-length, which has been demonstrated in other studies with potential long-term fitness implications (Mundinger, et al. 2021). This could indicate that the breeding females in our studied population to some degree buffered detrimental effects of poorer weather conditions on the postnatal development of their pups.

Colder temperatures in the early gestation period increased the chances of pregnant females leaving the colony; however, we observed that warmer temperatures in the late gestation period also had a negative effect on daily presence probabilities. In the early stages of pregnancy, we expect that females left the heated bat box for more suitable roosting conditions to enter torpor until weather conditions improved (Willis, et al. 2006). This was supported by our findings that absence during early gestation prolonged the gestation duration with equally many days. However, the observations of pregnant females leaving the box in response to high temperatures, despite the heating being turned off on such days, likely is a result of the roosting conditions reaching inhospitable levels in the colony (Bartonička and Řehák 2007). Absence in response to high temperatures did not prolong gestation duration, and one female gave birth while absent from the maternity roost before returning with her (estimated) 2-day old pup. Although such roost-switches can inflict extra costs on heavily pregnant females, highlighting the importance of proper bat box designs (Tillman, et al. 2021), a high latitude climate is perhaps not warm enough to result in artificial roosts becoming ecological traps, as was observed for bats in the Mediterranean (Flaquer, et al. 2014). Still, in light of ongoing climate change, the design and placement of bat boxes at high latitudes should also be considered given the potential lethal consequences of overheating (Crawford and O’Keefe 2021).

Across the seven breeding seasons a total of 28 viable pups were born, of which only seven females, resulting in a sex ratio of 3:1 in favour of male pups. Because the female pups were born over years to different mothers (see Table S1) we could not detect any apparent cause for this strong bias towards male pups. However, Barclay (2012) suggested that offspring sex-ratios in bats likely varies with the length of the breeding season, in addition to maternal qualities and environmental conditions. This was explained through reproductive success in one-year old females being more dependent than males on their juvenile body condition by the end of summer. Female pups should therefore be most beneficial to produce if born early in the season because it increases their chances of surviving the winter and returning as reproductive females to the colony (Frick, et al. 2010). No timing differences between genders were apparent in our data, but we observed that female pups expressed higher growth rates than male pups during the most rapid growth period, although we did not find a sex effect on the final size in juveniles reaching adult sizes. The elevated growth rates could suggest that female pups are more costly to produce, particularly when faced with the short breeding season at this high latitude, and will perhaps be a more careful investment than producing a male pup. However, because male pups disperse, we do not know survival rates in the males born to this study population, and such speculations will therefore remain untested.

We observed apparent adaptations to the short breeding seasons associated with a high latitude environment in the postnatal development of *E. nilssonii*: the pups were born big (with ∼41% of maternal forearm-lengths), grew fast (most rapid growth period occurring within 11 days), and became volant already after two weeks. Our observations of volant pups approaching adult flight patterns (emergence time and flight durations) by the end of their first flight week aligns with those made for *Myotis lucifugus* by Buchler (1980). However, our observations of pups flying for the first time as early as 13 days after birth supports the estimated timing of volancy in *E. nilssonii* reported by (Rydell 1989a, Rydell 1989b; earliest volancy recorded 12-17 days after hearing the first baby isolation calls), and is, to our knowledge, the earliest age of volancy recorded in any bat species to date.

### Conclusions

The unique day-to-day monitoring of individual-level breeding phenology and postnatal development in *E. nilssonii* presented in this study provides three main novel insights: firstly, we show evidence of strong and consistent maternal differences in breeding phenology in bats across years. This observation could indicate that reproductive female bats apply individual strategies across a behavioural continuum with two extremes; arriving early to prolong the breeding season but risking longer periods with poor foraging conditions that require temporal use of torpor; or arriving later when environmental conditions have improved but facing a shortened breeding season. However, these strategies are likely strongly connected with individual overwintering behaviour (Jonasson and Willis 2011, Reusch, et al. 2023) and should be further explored. Secondly, we present indications of reproductive females attempting to buffer detrimental environmental effects on the offspring development, by reducing the duration of gestation when arriving at the colony late (relative to own mean arrival dates) and by intensifying maternal investment to increase forearm-length growth rates in pups born late (relative to own mean parturition dates). Thirdly, we show that this population of *E. nilssonii* produce large, fast-growing pups with the age of volancy corresponding to that found in Swedish *E. nilssonii* colonies, which is the earliest recorded volancy currently observed in any bat species. Our observations offer potential insights to how *E. nilssonii* is adapted to reproducing in high latitude environments and thus potentially how it has become the northernmost breeding bat species in the world.

## Acknowledgements

We sincerely thank Keith Redford and Karl Kugelschafter (Chirotec) for assisting with the infra-red based surveillance of the bat box, Ronny Steen (NMBU) for financing and installing the video surveillance, Rune Sørås and Anke Kirkeby and several volunteers for helping out with providing mealworms, measurements and photo documentation, Erik Korsten for providing useful literature background, and we thank Thomas Lilley for corrections and comments on the manuscript. We also would like to acknowledge Jürgen Gebhard for being a great source of inspiration for this research. The study is dedicated to the memory of Jens Rydell (†).

