## Supplementary Materials for "Individual variation in breeding phenology and postnatal development in northern bats (*Eptesicus nilssonii*)"

### Sex ratio of pups per mother and year

Throughout the 7 breeding seasons, a total of 7 female pups and 21 male pups were born to the colony (in addition to one still-born Siamese twin). An overview of the sex-ratio per mother and year is shown in Table S1.

**Table S1**: Overview of the sexes of pups born to each female per year. Years written in the header indicates birth-year of each mother. Females marked with an asterix (*) were hand-raised as pups. Bat9 is included, although she was unreproductive in her first year back in the box, which was the last year of the study.

| | Bat1* (2015) | Bat2* (2015) | Bat3* (2017) | Bat4* (2017) | Bat5 (2017) | Bat6* (2017) | Bat7* (2017) | Bat8 (2020) | Bat9 (2022) |
|---|---|---|---|---|---|---|---|---|---|
| 2017 | M | F (Bat5) | | | | | | | |
| 2018 | M | M | M | M | Siamese | | | | |
| 2019 | M | M | | M | F | M | M | | |
| 2020 | F (Bat8) | M | | M | F | | | | |
| 2021 | | M | | M | F | | | M | |
| 2022 | | F (Bat9) | | M | M | | | M | |
| 2023 | | M | | M | M | | | F | Unrep. |

### Umbilical cord and placenta

Figure S1 shows pictures taken shortly after one of the births, where the umbilical cord and placenta were still attached to the newborn pup.

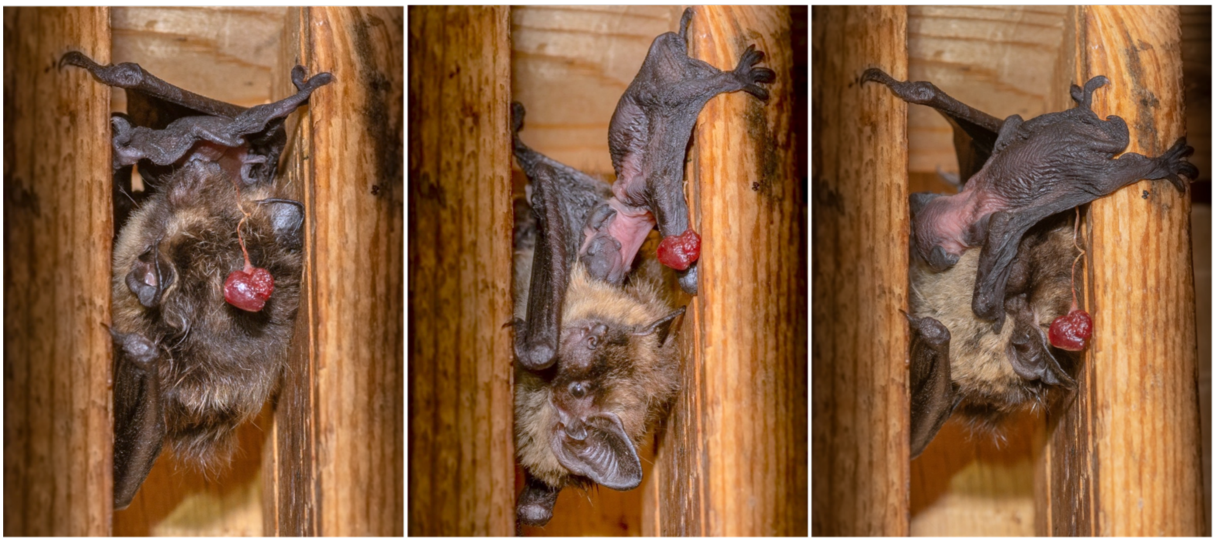

**Figure S1**: Three photos of a female nursing her newborn pup, with the placenta still being attached to the umbilical cord. Photos by Jeroen van der Kooij.

**Size at birth**

Effects from the best model explaining the variation in forearm-length at birth are shown in Figure S2.

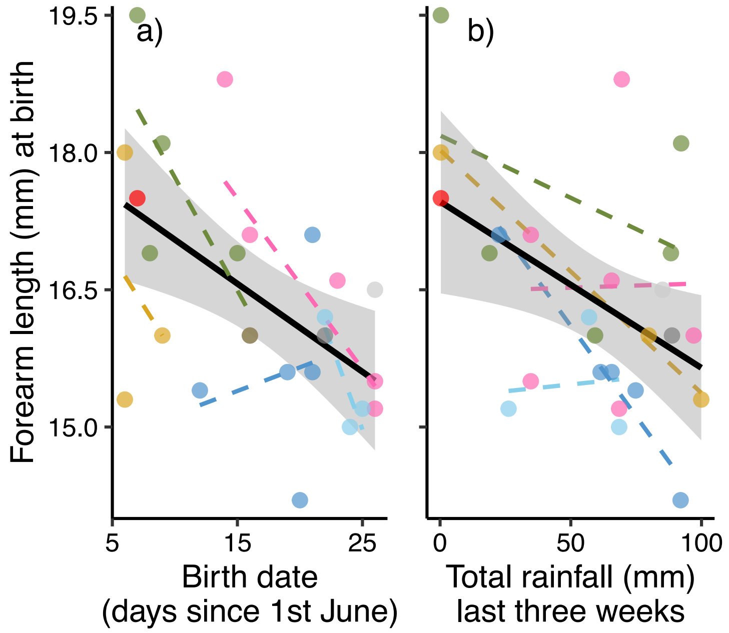


**Figure S2**: Effects on forearm length at birth, including **a)** a negative effect of later birth dates and **b)** a negative effect of wetter (and colder) weather conditions the three weeks prior to parturition. The black regression lines show the overall original models effects while different colours indicate different mothers.

**Growth models and individual growth patterns**

Parameters from the three models tested on the postnatal growth are shown in table S2. Individual growth of forearm length (Fig. S3) and body mass (Fig. S4) are illustrated against the best fitted growth models (logistic for forearm length and von Bertalanffy for body mass).

**Table S2**: Growth parameters derived from the logistic, Gompertz, and von Bertalanffy growth models. The parameters are: *A* = asymptotic value, *K* = growth rate constant, *I* = inflection point. The models were fitted based on respectively 450 measures of forearm length and 424 measures of body mass from 26 bat pups, excluding one pup with unknown birth-time and one pup that died 8 days old.

| Growth model | Parameter | Forearm length across age (days) | | | Body mass across age (days) | | |
|---|---|---|---|---|---|---|---|
| | | Estimate | SE | ΔAICc | Estimate | SE | ΔAICc |
| Logistic | *A* | 39.94 | 0.158 | 0.0 | 8.40 | 0.09 | 5.3 |
| | *K* | 0.25 | 0.005 | | 0.27 | 0.015 | |
| | *I* | 1.92 | 0.059 | | 2.91 | 0.163 | |
| Gompertz | *A* | 40.39 | 0.187 | 47.6 | 8.53 | 0.104 | 0.2 |
| | *K* | 0.20 | 0.004 | | 0.20 | 0.012 | |
| | *I* | 0.09 | 0.065 | | 1.14 | 0.170 | |
| von Bertalanffy | *A* | 41.04 | 0.240 | 130.0 | 8.74 | 0.120 | 0.0 |
| | *K* | 0.15 | 0.004 | | 0.14 | 0.009 | |
| | *I* | -2.80 | 0.116 | | -1.86 | 0.253 | |

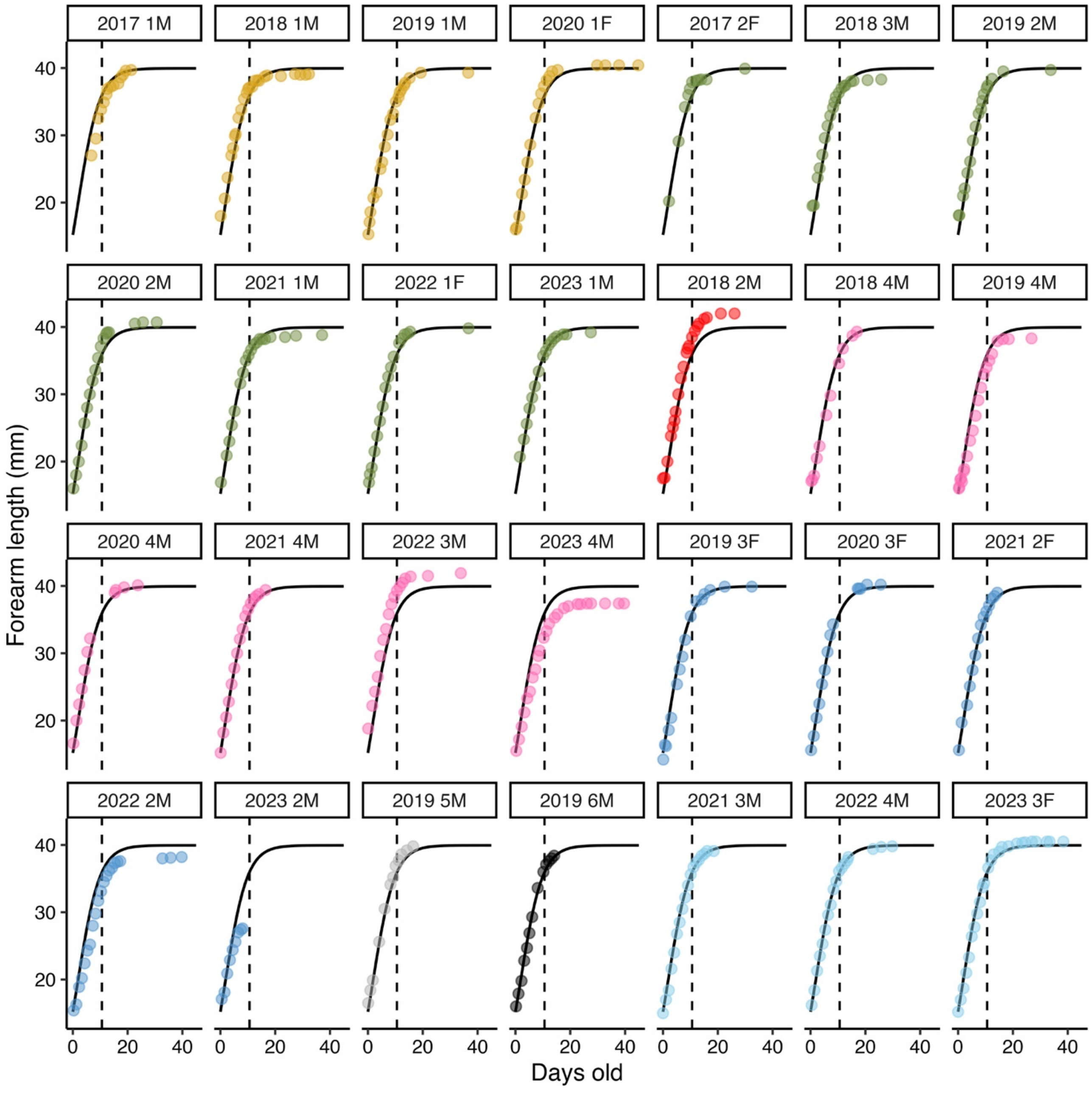


**Figure S3**: Individual forearm length growth for each pup born in the colony. Different colours indicate different mothers (same colour-coding as in the main manuscript), with datapoints being the measurements for each pup, and the solid growth curve in each window showing the best fitted logistic model based on all datapoints, for comparison. The dashed vertical line in each window indicates the break-point at 10.6 days for the most rapid growth period. The name of each window corresponds to the year of birth, along with the order of which the pup was born in that year, and finally the sex of the pup (M = male, F = female). Pup '2023 2M' was the male pup that died of unknown causes when he was 8 days old, and pup '2019 3F' was the female pup that did not return from her first flight when she was 14 days old.

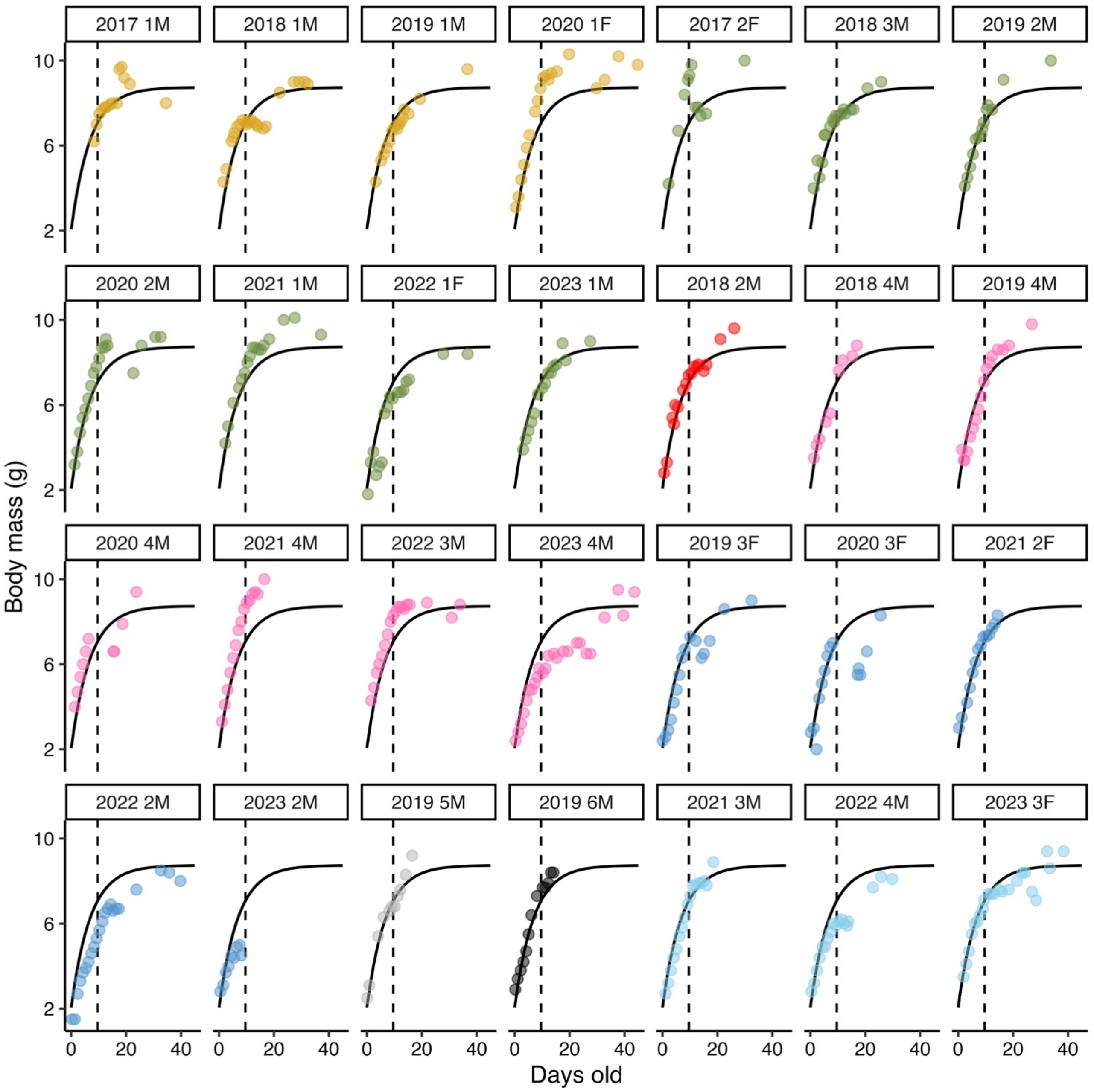


**Figure S4**: Individual body mass growth for each pup born in the colony. Different colours indicate different mothers (same colour-coding as in the main manuscript), with datapoints being the measurements for each pup, and the solid growth curve in each window showing the best fitted von Bertalanffy growth model based on all datapoints, for comparison. The dashed vertical line in each window indicates the break-point at 9.6 days for the most rapid growth period. The name of each window corresponds to the year of birth, along with the order of which the pup was born in that year, and finally the sex of the pup (M = male, F = female).

**Juvenile flight patterns**

Details for each flight night during juveniles' first flight week are shown in Table S3 and Table S4.

**Table S3**: Emergence time (i.e. minutes since sunset) for juveniles and adults on the juveniles' first flight week (flight night 1 = the night of the juveniles' first flight).

| | Juveniles | | | | Adults | | | |
|---|---|---|---|---|---|---|---|---|
| Flight night | Min | Max | Mean (± SD) | $N_{obs}$ | Min | Max | Mean (± SD) | $N_{obs}$ |
| 1 | 48 | 275 | 162 ± 73.6 | 10 | 0 | 100 | 28.5 ± 18.6 | 38 |
| 2 | 27 | 147 | 106 ± 44.1 | 10 | -1 | 100 | 24.5 ± 18.9 | 39 |
| 3 | 35 | 101 | 54.9 ± 19.8 | 8 | 0 | 50 | 30.3 ± 12.1 | 29 |
| 4 | 35 | 140 | 53.8 ± 35.2 | 8 | 11 | 49 | 31.6 ± 9.6 | 29 |
| 5 | 34 | 53 | 42.6 ± 6.9 | 7 | 20 | 45 | 30.4 ± 8.0 | 25 |
| 6 | 23 | 41 | 33.7 ± 8.0 | 10 | 3 | 52 | 25.1 ± 10.5 | 33 |
| 7 | 24 | 46 | 36.5 ± 8.4 | 6 | 12 | 46 | 29.8 ± 9.9 | 23 |
| 8 | 13 | 45 | 28 ± 12.3 | 6 | -16 | 55 | 22.6 ± 17.2 | 25 |

**Table S4**: Duration time (minutes) of trips registered for juveniles and adults on the juveniles' first flight week.

| | Juveniles | | | | Adults | | | |
|---|---|---|---|---|---|---|---|---|
| Flight night | Min | Max | Mean (± SD) | $N_{obs}$ | Min | Max | Mean (± SD) | $N_{obs}$ |
| 1 | 1.3 | 40.6 | 15.3 ± 15.8 | 7 | 5 | 252 | 62.7 ± 45.2 | 46 |
| 2 | 14 | 72 | 35.4 ± 22.7 | 9 | 5.1 | 250 | 81.9 ± 58.3 | 46 |
| 3 | 11.7 | 65.4 | 42.4 ± 23 | 5 | 18.9 | 201 | 101 ± 50.2 | 18 |
| 4 | 9.1 | 121 | 45.8 ± 38.7 | 9 | 8.2 | 182 | 95.7 ± 59.8 | 26 |
| 5 | 5.9 | 126 | 54.3 ± 42.8 | 8 | 16 | 235 | 107 ± 57.4 | 23 |
| 6 | 2 | 100 | 37.1 ± 34.9 | 9 | 3.2 | 184 | 63.6 ± 41.9 | 30 |
| 7 | 20 | 135 | 80.7 ± 46.8 | 6 | 7 | 237 | 111 ± 59.2 | 24 |
| 8 | 22.2 | 176 | 101 ± 70.9 | 5 | 40.2 | 210 | 113 ± 60.3 | 14 |
